# Predictive visual uncertainty around moving trajectories influences causality judgments in launching displays

**DOI:** 10.64898/2026.02.06.704483

**Authors:** Lina Eicke-Kanani, Fabian Tatai, Lukas Rosenberger, Christina Schmitter, Benjamin Straube, Thomas S. A. Wallis

## Abstract

Michotte’s “launching displays” are animations of collision-like interactions between two objects that elicit a stable and robust impression that one object, the launcher, caused another object, the target, to move. Although it is well-known that unexpected disruptions of movement continuation between launcher and target decrease causal impressions in centre-to-centre collisions, the role of observers’ visual uncertainty around predicted moving trajectories remains relatively unexplored. In this work, we (1) assess observers’ uncertainty around post-collision moving angles in a trajectory prediction task and (2) collect their causal impression in a causality rating task. In the latter task, observers viewed centre-to-centre collisions with different levels of movement continuity between the launcher and the target disc. By presenting different launch orientations, we exploited the well-known oblique effect to vary trajectory prediction uncertainty within individuals. If observers rely on their trajectory predictions to rate the causality of the collision, we expect their accuracy in (1) to have a systematic influence on their causality rating in (2). We replicate previous findings that observers report stronger causal impressions in trials where the target and the launcher move in the same direction and weaker causal impressions for collisions where the target and the launcher moving trajectory deviated. Furthermore, causality ratings were on average higher for oblique compared to cardinal launch directions, implying that increased sensory uncertainty induces a stronger causal impression. We hope this work will inspire deeper empirical assessments and computational models describing the role of sensory uncertainty and predictive processes in shaping subjective impressions of causality.

## Introduction

In his seminal work, Michotte (1963) demonstrated that observers tend to perceive a physical collision if a 2D object abruptly moves after being contacted by another 2D object. Observers viewing such launching displays frequently report the perception of a physical collision where the first moving object, the launcher, was the cause of the second object’s movement, the target (Michotte, 1963; White, 2017). The subjective impression that one object launched another into motion can be regarded as a subjective impression of physical causality or an *efficient cause* according to Aristotle (Falcon, 2006). In an efficient cause, the observer is concerned with the question of whether the second object’s motion stemmed from its collision with the first object.

Michotte (1963) concluded that even early stages of visual processing could detect causal relationships, due the robustness of causality judgments in response to different launching displays. If causality is indeed encoded at early stages of visual processing and therefore constitutes a perceptual domain (see for example Kominsky & Scholl, 2020; Ohl & Rolfs, 2023; Rolfs et al., 2013; though see Arnold et al., 2015), causality detectors should be susceptible to systematic changes in an observer’s visual uncertainty such as uncertainty around movement direction. The view that causality perception should be subject to visual uncertainty parallels ideas from intuitive physics research, where the causality judgment is believed to reflect an observer’s internalised knowledge of Newtonian physics under uncertainty (Sanborn et al., 2013).

The importance of the launcher’s movement direction for a subjective causal impression – namely the notion that the launcher is the efficient cause for the target’s motion – has been highlighted by evidence on retinotopic adaptation to launch direction (most notably by Ohl & Rolfs, 2023; also see Kominsky & Scholl, 2020 and Rolfs et al., 2013), as well as intuitive physics work (Gerstenberg et al., 2012, 2014, 2021; Smith & Vul, 2013) and research on causality judgments itself (Schülke et al., 2023; Straube & Chatterjee, 2010; White, 2012). The latter studies provide evidence that causal impressions are reported more frequently if the launcher and the target have the same moving direction after colliding head-on; in contrast, launching displays are judged as less causal the more the launcher’s moving direction and the target’s moving direction deviate from one another after a centre-to-centre collision (Schülke et al., 2023; Straube & Chatterjee, 2010; White, 2012). Since in a real-world centre-to-centre collision with a rolling movement, the target trajectory should be a continuation of the launcher’s trajectory, the following text will use the words “plausible” and “plausibility” to express that the launcher’s and target’s moving directions were similar.

The fact that the subjective causality judgment decreases in centre-to-centre collisions when the target trajectory disrupts the movement continuation of the launcher’s trajectory (Schülke et al., 2023; Straube & Chatterjee, 2010; White, 2012) indicates a systematic relationship between the real-world implications of physics and an observers’ causal impression (Sanborn et al., 2013). Intuitive physics frameworks have emphasised the role of individuals’ uncertainty, namely, that real-world physical properties are inferred under individual noise given an internal physics model (Smith & Vul, 2013). Furthermore, intuitive physics work has highlighted the importance of plausibility-based noisy mechanisms to predict future outcomes (Gerstenberg et al., 2012, 2014, 2021; Smith & Vul, 2013). The idea of plausibility-based mechanisms usually assumes an internal model on the observer’s side that is aligned with the real-world implications of physics. Additionally, these so-called “Noisy Newtonian” frameworks consider the observer’s perceptual uncertainty around physical quantities in the perception of causality (Sanborn et al., 2013). However, that same line of research does not assess individual observer’s uncertainty empirically, and average causality reports across subjects are explained with general model parameters. This makes it difficult to evaluate the exact link between perceptual uncertainty and causality judgments.

Therefore, to allow us to relate causal impressions and sensory uncertainty, we aimed to 1. empirically measure the accuracy of observers’ predictions around the target’s moving trajectory after contact with the launcher and 2. collect causality ratings from the same observers on a slider scale in launching displays with plausible and implausible target moving trajectories. In the latter task, the target movement trajectory was manipulated by systematically adding an offset to the target’s movement continuation of the launcher’s trajectory. Our research question was the following: Do observers rely on their uncertainty around the target’s moving trajectory to judge causality in launching displays?

If the real-world causality of a collision is indeed detected at early perceptual stages and informs the causality judgment, then visual uncertainty (uncertainty around movement trajectories) should systematically affect causal impressions; the same is true if the causality judgment is generated from inferences about the physics of the collision: if causality judgments are the result of a plausibility-based mechanism, then there should be a clear relationship between visual uncertainty (i. e. uncertainty around predictions of moving directions) and causal impressions.

For instance, imagine viewing a centre-to-centre collision between two balls, where the launcher approaches the target (Figure 1, left, pre-collision) which then also moves in the same direction with a similar speed (Figure 1, right, post-collision). If the objects move at an appropriate speed, it should be clear to the observer that the collision happened head-on and that the target should move in the same direction as the launcher. Under a model where the target and the launcher should move in the same direction, even with perceptual uncertainty, the causal impression for a plausible collision should be strong because the target’s observed trajectory is similar to the target’s predicted trajectory. In contrast, the more the target motion violates the observer’s predicted trajectory, the more the causal impression should decline.

**Figure 1.**
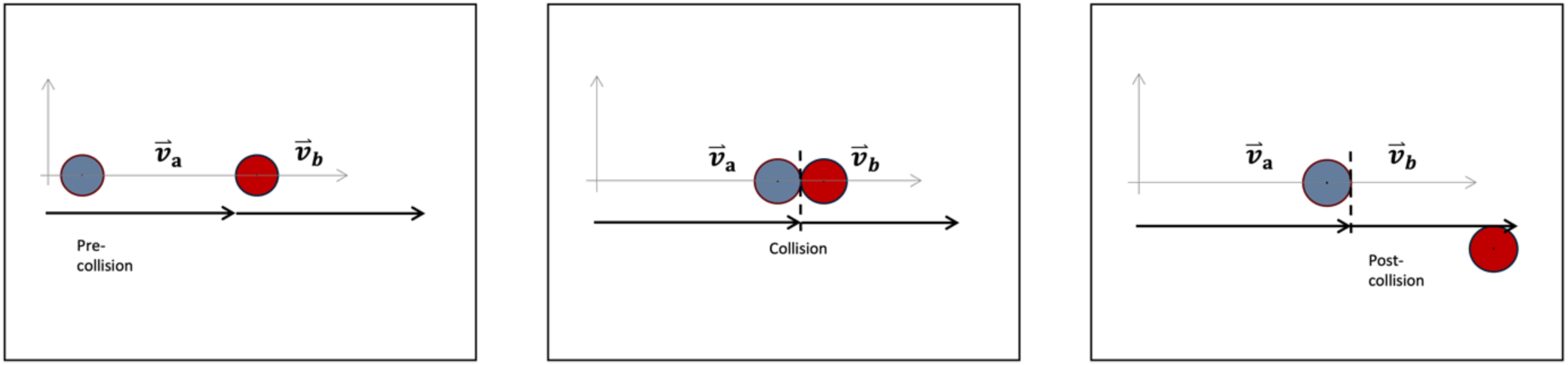
Physically plausible moving trajectories are easily predictable on a horizontal plane. The launcher’s (blue) velocity vector before the collision (left) informs an observer’s prediction of the outgoing target’s trajectory (red) after collision, which we assessed in a trajectory prediction task at the collision time (middle). In a second task we assessed subjective causality in response to the target’s moving trajectory which could be contradicting the prediction as in this example (right); we expect implausible trajectories to decrease observer’s causal impression. It has been proposed that observers rely on noisy internal simulations to inform predictions of moving trajectories and outcomes to judge causality (Gerstenberg et al., 2021; Sanborn et al., 2013; Smith & Vul, 2013)

For the assessment of sensory uncertainty, observers viewed launching displays up to the point of collision and then indicated their prediction of the target’s end position after the collision in an adjustment task. Afterwards, the same observers viewed full launching displays (pre- and post-collision state) and judged the causality of the collision on a slider scale. In a previous study, we already showed that changing the orientation of the launch can lead to changes in visual uncertainty around the collision point (more errors in reported collision point for oblique launches than for cardinal launches) and to systematic changes in causality reports (Duan et al., 2025). In that study, we assumed that the collision point is the key frame that will determine an observer’s prediction of the target’s moving trajectory after collision.

In the current work, we aim to extend that line of reasoning by measuring the observer’s actual predictions of moving trajectory for different launch directions directly. Varying the orientation of the collision should vary the visual uncertainty in both, the prediction and the causality judgment task, which could allow us to investigate causal impressions under different degrees of visual uncertainty. Under the assumption that observers rely on their uncertainty around moving trajectories to simulate post-collision outcomes, their uncertainty should be directly related to how observers judge causality.

A possible caveat in assuming a plausibility-based mechanism underlying causality judgments is the inherent subjectivity of the causality judgment that may not be fully explained by uncertainty around the launcher’s and target’s movement direction or internal noise in the observer’s process model: causality judgments require a decision component and can be evoked even for scenarios that are physically impossible (Bechlivanidis et al., 2019, 2022). Moreover, observers report causality even for abstract launcher-target-interactions, i. e. colour transitions upon collision between the launcher and target object or social causality (Blos et al., 2012; Kruschke & Fragassi, 2019; Wende et al., 2013, 2015), implying that a subjective decision process could be foundational to behavioural causality judgments.

## Methods

### Participants

35 participants (17 male, 18 female) with a mean age of 24.4 years (SD=3.2 years) participated in behavioural experiments and gave their informed consent. Experiments were approved by the ethics committee of the Technical University of Darmstadt (EK56/2022), and conformed to Standard 8 of the American Psychological Association’s Ethical Principles of Psychologists and Code of Conduct (2017) and to the Declaration of Helsinki (with the exception of Article 35 concerning preregistration in a public database). Participants were either volunteers who did not receive any compensation or received course credit.

Participants had normal or corrected-to-normal vision. We consider participants neurotypical, meaning they were inside the norm based on the results of the O-LIFE schizotypy questionnaire (Mason et al., 2005); we considered that information important as schizophrenia has been associated with altered perceptions of causality (Schülke et al., 2023; Wende et al., 2015).

### Equipment and Software

Experiments were programmed in Python 3.6 under Ubuntu 20.04. using PsychoPy 2022.1.4 and pypixxlib 3.12.12226 for rendering on a PROPixx projector controlled via the DATAPixx 3 (VPixx Technologies, Saint-Bruno-de-Montarville, Canada). The image was projected onto a 0.97 m by 0.55 m screen. Participants sat in a chin-rest 130 cm away from the screen, resulting in a display of 41 x 24 dva and a launch distance travelled by the discs of 21.6 dva. The launcher disc and the target disc had a radius of 0.9 dva.

The physics of the collision was simulated using the physics engine pooltool version 0.1 (Kiefl, 2024); we assumed a frictionless rigid collision between two rolling objects (no spin) and extracted the object positions on each timeframe by passing pooltool the distance between the objects, the speed and the desired key frame of the collision (midpoint of the distance).

### Stimuli and Procedure

All participants performed a trajectory prediction task followed by a causality rating task. Responses from the prediction task were used to compute observer’s sensory uncertainty around the target ball’s moving direction after the collision. The causality rating task was then conducted to assess individuals’ subjective causal impressions in launching displays where the target ball’s exhibited movement directions was systematically rotated from the launcher ball’s moving direction.

In the trajectory prediction task, participants were instructed to report the position where they believed the target ball should be headed after the collision. On each trial, the angle of collision between launcher and target ball (Figure 2A) was randomly sampled from a uniform circle. In both of our tasks, launching positions were initialised randomly around a virtual (invisible) circle with a diameter of 2.1 dva to avoid local adaptation to the dynamics of the collision (Ohl & Rolfs, 2023; Rolfs et al., 2013). At the beginning of each trial, observers saw the launcher’s and target’s start position for one second before the launcher started to move (speed was uniformly sampled between 11.9 to 36.4 dva/sec) until touching the target.

**Figure 2.**
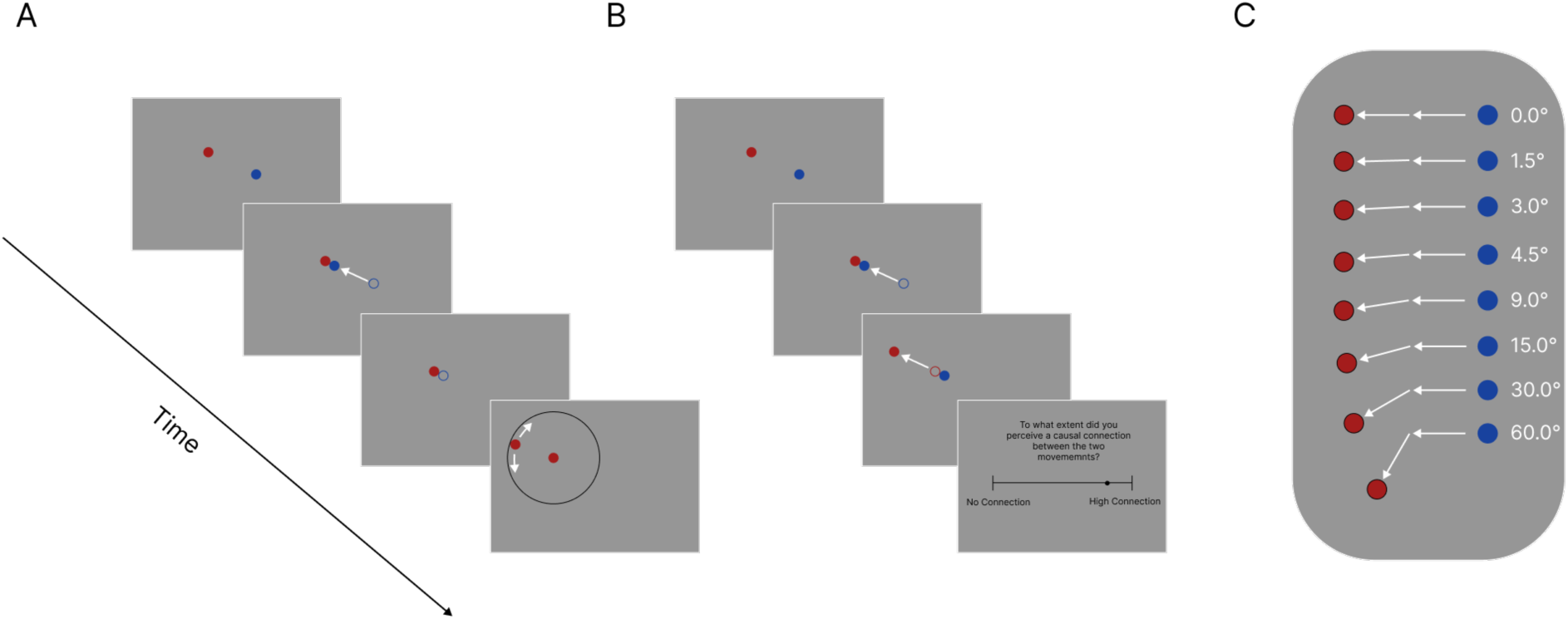
Experimental Paradigm. **A.** Participants performed a prediction of moving trajectory task where they only viewed the launcher ball motion until the point of collision with the target. Afterwards, they indicated the outgoing moving direction the target should have in an adjustment task. **B.** Participants performed a causality task where they viewed a full launching display (target and launcher movement). After the launcher collided with the target, the target either moved in the same direction as the launcher or disrupted the movement continuation by moving with a relative angle offset. Afterwards, participants rated for each interaction how connected the launcher and the target movement seemed to them (from no connection to high connection). **C.** The target offsets employed to violate the continuation of the launcher’s movement in B were sampled from a deterministic list comprising +/- [0.0, 1.5, 3.0, 4.5, 9.0, 15.0, 30.0, 60.0]°, where 0.0° means no violation of movement continuation and 60° means high violation of movement continuation.

Participants always performed a block of 420-500 prediction trials followed by a block of 400-480 causality trials (depending on participant’s energy and focus which they communicated to the experimenter). Participants were allowed to take a short break every 10 minutes and a longer break between the experimental blocks. The total experiment time for both tasks ranged between 90-120 minutes.

In the prediction task, participants only saw the pre-collision and the collision phase (Figure 1); on the response screen. The launcher disappeared upon touching the target so that participants could not rely on the launcher’s end point to extrapolate the target’s moving direction. As a tool to give the response in the adjustment task, another disc that looked identical to the target disc appeared on the screen at a fixed distance of 8.5 dva from the target’s position on the collision frame. Participants adjusted the position of that target ‘twin’ using the computer mouse and clicked the mouse button to indicate their prediction of the target’s end position (Figure 2A, see <u>here</u> for a demo).

Prior to the causality judgment task, participants performed a practice session viewing maximally plausible (perfect launcher movement continuation) and maximally implausible (60° offset from launcher movement continuation) target moving trajectories and rating their causality. The purpose was to prepare participants for the task by showing them extreme movement trajectories and making them familiar with the slider scale to report causality. Participants learned the task intuitively in the practice session and received no further instructions or additional information about the collision.

In the causality judgment task (Figure 2B), each trial was initially similar to the prediction task until the collision frame. At the collision time, when the launcher stopped and the target started moving, participants either viewed a physically plausible trajectory (no violation of continuity) or an implausible trajectory (violation of continuity = angle offset of launcher movement direction, see Figure 2C). Launcher and target always made direct contact and the target always moved immediately after being contacted by the launcher without any delay.

The target’s moving trajectory was determined relative to the launcher’s moving trajectory: in a physically plausible case of a centre-to-centre collision, the target should always move in the same direction as the launcher. The target’s trajectory followed a plausible scenario (0° offset from the launcher’s trajectory, Figure 2C, top) in 50% of the trials. In the rest of the trials, the target’s moving trajectory relative to the launcher’s moving trajectory was sampled uniformly from a list of angle offsets +/- [1.5, 3.0, 4.5, 9.0, 15.0, 30.0, 60.0]°. Note, that the +/- 9° offset was only employed for 12 of the participants to cover the range between +/- 4.5 and +/- 15.0 after looking at preliminary data. On the population level, a minimum of 480 trials (9°) and a maximum of 9580 trials (0°) were performed per angle offset. The offsets are signed because they can be added as a clockwise or counterclockwise manipulation of the launcher’s movement.

At the end of each causality trial, observers answered the following question on the response screen: “To what extent did you perceive a causal connection between the two movements?” on a slider ranging from “no connection” (left end of the slider) to “high connection” (right end of the slider). The response item on the slider was always centred at 0.5. Participants indicated their causal impression by moving the slider with the mouse to the right (closer to 1.0, high connection (causal)) or the left (closer to 0.0, no connection (not causal)) and confirmed their response by a key press.

### Data Analysis

Data analysis was performed using pandas, numpy and scipy.stats with visualisations in seaborn and matplotlib under Python 3.8.16. Circular statistics were implemented after Jammaladaka and Sengupta (2001).

#### Circular statistics

Circular computations of descriptive statistics generally follow the logic that direction can be expressed as a vector and consequently, the centre of a circular distribution with a sample size *n* can be computed by computing a resultant vector ***R*** of all values defined as

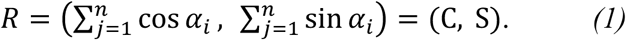

*α* denotes the single values in radians that are projected from a polar onto a rectangular plane (x- and y-direction) by taking the sum of the sine and the cosine, respectively (Jammalamadaka & Sengupta, 2001); C is the resultant vector in the y direction and S is the resultant vector in the x direction. The circular mean 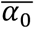 is then computed as

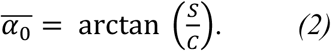

Under a similar logic, circular dispersion was computed as the difference in resultant vector length between *n* unit vectors with the same direction (a reference with length equal to *n*) and the resultant vector length ***R*** of the actual values in our dataset, assuming that each of them is also a unit vector with length 1. ***R*** can again be computed as in (1) or simplified according to Pythagoras’s theorem:

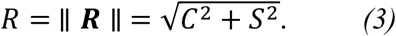

To show how dispersed (or concentrated) our response data is around a mean direction, it is sufficient to simply take the difference between the number of samples *n* (resultant vector length if all clicks had the same direction and unit length) and the empirical resultant vector length *R* of the clicks given by (1) and (3) to compute circular dispersion *D_v_*:

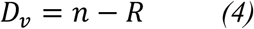

In addition, to scale the values for circular dispersion between 0 and 1 we divided them by the number of samples *n*. Therefore, values of 0 indicate the absence of dispersion (all trajectory responses are in the same direction) whereas a value of 1 would indicate circularly uniform responses.

Although the underlying data was circular, the circular dispersion is a linear (albeit bounded) value; therefore, we used the arithmetic mean to compute the group-level dispersion by taking the mean of individual participants’ dispersion value per experimental condition. The circular mean in contrast remains circular and we computed grand means by taking the circular mean once more.

#### Data aggregation

Data was aggregated over dichotomized launch directions, cardinal and oblique, under different binnings of +/- 2.5°, 5.0° and 7.5° around exactly cardinal (0°, 90°, 180°, 270°) and exactly oblique (45°, 135°, 225°, 315°) moving angles (separately for the launcher and target moving angle see text below). Taking the mean per participant (arithmetic for causality data and circular for prediction data from the adjustment task) over those bins allowed us to perform a paired Wilcoxon signed-rank test to quantify cardinal-oblique differences. In the prediction task, the circular mean and the circular dispersion were computed per participant and for cardinal and oblique launches, serving as input data for the Wilcoxon signed-rank test. From that data, we also computed grand averages of the prediction errors and of the circular dispersion; the latter served as a measure of sensory uncertainty (see (4)).

As our initial hypothesis was that the launcher’s movement before collision should determine the prediction of the target’s post-collision moving direction, we expected to find that both the prediction accuracy of the target ball’s end position and the mean causal impressions should be altered in cardinal launcher moving directions vs. oblique launcher moving directions. In the causality judgment task, both the pre- and the post-collision phase were displayed, meaning that we applied the binning for the cardinal / oblique classification separately to the launcher’s moving angle and the target’s moving trajectory. Causal impressions per participant were computed separately per participant in cardinal / oblique launcher trajectories and also in cardinal / oblique target trajectories to perform a Wilcoxon signed-rank test as in the prediction data.

To address the role of the plausibility of the target moving angle, we identified post-hoc when the violation of movement continuation in the target’s moving trajectory disrupted causal impressions (mean causal impressions dropped below 0.5). We found that such a change in causal impressions happened for angle offsets > 15.0° which is consistent with White (2012) who reported that causal impressions become significantly weaker for an angle offset of 20°.

That categorisation was used to define a ‘high continuity’ and ‘low continuity’ label. In a 2 x 2 analysis design, we computed a single mean causal impression per participant for high continuity vs. low continuity moving angles and for cardinal vs. oblique launcher moving trajectories. Lastly, we aimed to correlate the results from the causality task with the results of the prediction task; to do so, we computed a regression line using the single-participant sensory uncertainty (computed as the circular dispersion) in cardinal vs. oblique launcher moving trajectories and the mean causal impressions in high continuity vs. low continuity trials under cardinal vs. oblique launcher moving angles. Note, that the prediction task did not have a post-collision phase, meaning that the same cardinal / oblique uncertainty was correlated separately with the mean causal impressions for high continuity and low continuity trials per launcher moving angle, respectively. The regression was computed in np.polyfit and the spearman correlation was computed in scipy.stats.spearmanr.

## Results

In the prediction task, participants viewed a launching display up to the point of collision between the launcher and the target disc. Afterwards, the target disappeared and participants indicated the target’s predicted end position in an adjustment task. Participants on average performed the task with good precision (Fig. 3A). Specifically, we computed the angular difference between the physically plausible moving angle (equal to the launcher’s moving angle) and the participants’ click location on each trial (displayed vs. reported). The resulting histogram is centred around 0 (*M*=0.03°, *SD*=10.43°), meaning that participants’ predictions were firmly aligned with the physical ground truth.

**Figure 3.**
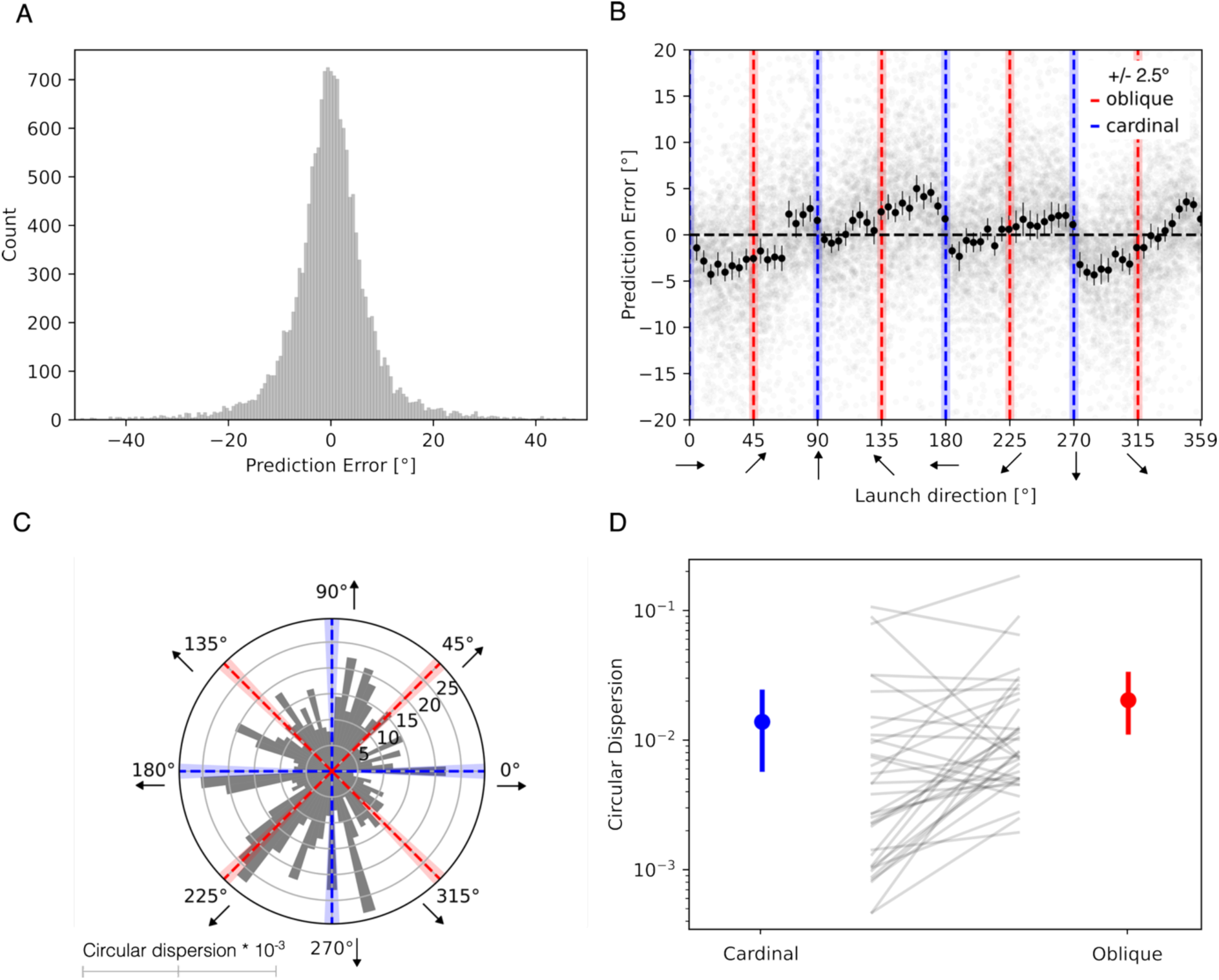
Uncertainty around predictions systematically depends on launch direction. **A.** Frequency of report errors (plausible trajectory minus reported trajectory) in a prediction of moving trajectory task across all participants. **B.** Biases in the prediction task systematically depend on launch orientation. Faint gray points show individual trial prediction errors, namely the angular distance between their prediction and the plausible (continuation of the launcher ball’s trajectory) target moving trajectory. Black dots show the mean bias (mean of participant means) within each +/- 2.5° bin of launch direction with error bars depicting the standard error of the mean. **C.** Sensory uncertainty around the prediction varies depending on launch direction. The grey bars on the polar plot depict the mean of participants’ mean uncertainty within each +/- 2.5° bin of launch direction; uncertainty was computed as the circular dispersion of the prediction errors in a bin. Exactly cardinal and exactly oblique locations +/- 2.5 degree bins are marked in blue and red, respectively. Those locations are the foundation for the mean uncertainties depicted in D. **D.** Sensory uncertainty in cardinal launches was on average lower than sensory uncertainty in oblique launches. The dots depict the mean uncertainty (arithmetic mean of individuals’ circular dispersions) and the 95% confidence interval (errorbar) for cardinal (blue) and oblique launches (red), respectively. The p-value is the result of a Wilcoxon signed-rank test.

Consistent with previous reports (Gros et al., 1998), we find that participants’ bias around the predictions depends on the launcher’s moving direction (Figure 3B). The mean bias, as measured by the circular grand mean of participants’ circular mean report errors per 5° bin of the launch direction (black dots with standard errors), is near zero at both cardinal and oblique orientations (Figure 3B, blue and red dashed lines) and varies systematically for intermediate launch angles. While bias is related to sensory uncertainty, additional assumptions and computations are required to transform between the two (Wei & Stocker, 2017) and we therefore do not use bias as an uncertainty measure in the present analysis.

To quantify participants’ uncertainty around predictions we computed the circular dispersion of report errors in 5° segments of the launch direction, following the logic that participants’ reports should be more variable when sensory evidence is weaker (Wei & Stocker, 2017). We then computed the average uncertainty across individuals by taking the arithmetic mean of these individual circular dispersions. Greater perceptual uncertainty in a bin should produce a higher average circular dispersion than in a bin with reduced perceptual uncertainty. The circular dispersion is shown as bars in the polar plot in Figure 3C.

Participants’ uncertainties depended strongly on the launch direction, but this relationship was highly variable (Figure 3C). For example, while some cardinal launch trajectories showed the expected lower dispersions (90° and 180°), others show comparatively high dispersions (0° and 270°). This pattern of results may indicate additional biases depending on the specific launch direction.

Nevertheless, when averaging over all cardinal and all oblique launches, we do observe the expected reduction in uncertainty for cardinal compared to oblique launches (Figure 3D). The average circular dispersions (blue and red dots with confidence intervals for cardinal and oblique launches, respectively) are shown in Figure 3D. Uncertainty was overall increased in oblique (red, *M=20*10^-3^*, *SE=5*10^-3^*) vs. cardinal (blue, *M=14*10^-3^*, *SE=4*10^-3^*) launches in +/-2.5° bins (Wilcoxon sign-rank *W=163*, *p=0.01*). Because the choice of +/- 2.5° bins was arbitrary, we repeated this analysis with bins of +/- 5.0° (*W=178*, *p=0.02*) and +/- 7.5° (*W=200*, *p=0.06*). We also tested the difference between cardinal trials vs. all other orientations, which yielded Wilcoxon sign-rank p-values of *p=2*10^-4^, p=0.03* and *p=0.11* for +/- 2.5, 5.0, 7.5° around the cardinals, respectively.

In the causality judgment task, observers viewed a full launching display where one disc (launcher) contacted a second disc (target) which moved after contact. The collision was always central and had neither spin nor friction, which meant that in a physically plausible scenario of a rigid two ball collision, the launcher and the target should move in the same direction. After collision, the target either moved in the same direction as the launcher (0° offset) or with an offset of +/- [1.5, 3.0, 4.5, 9.0, 15.0, 30.0, 60.0]° (see Figure 2C).

Figure 4 A-C shows the lowess for causal impressions in cardinal vs. oblique launches (blue and red lines) together with the raw response data (blue and red dots) per angle offset of the target’s moving trajectory from the launcher’s moving trajectory. Like in the prediction data, we tested different binnings for our cardinal / oblique classification of +/- 2.5°, 5.0° and 7.5°. Aligned with previous studies (Schülke et al., 2023; Straube & Chatterjee, 2010; White, 2012), we found that observers qualitatively report weaker causal impressions the higher the difference between the target’s and launcher’s moving directions.

**Figure 4.**
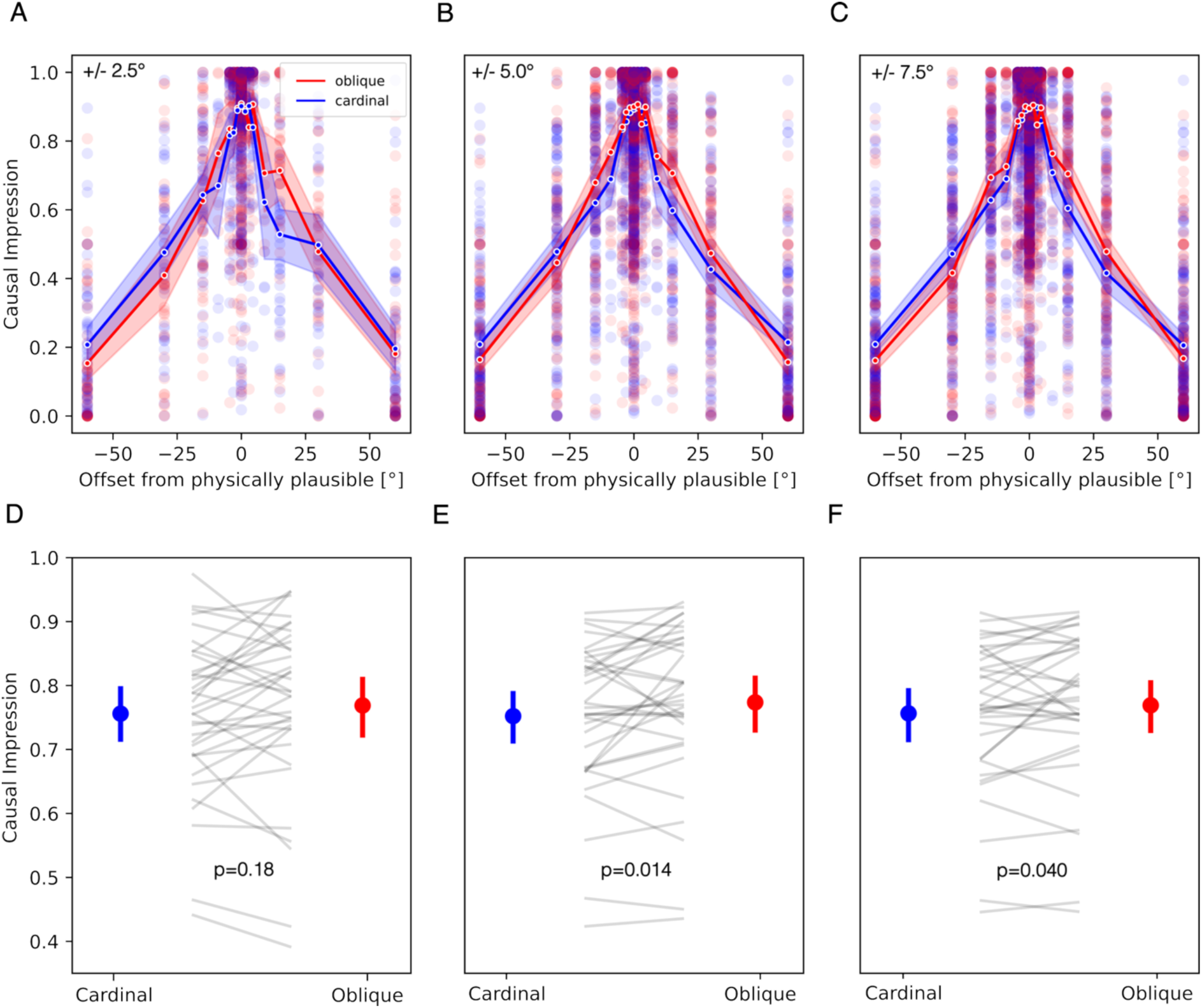
Causal impressions are influenced by the target’s relative movement continuation of the launcher’s cardinal vs. oblique trajectory. **A-C.** Causal impressions systematically changed with disruptions of movement continuation. The top row shows causal impressions for +/- [0.0, 1.5, 3.0, 4.5, 9.0, 15.0, 30.0, 60.0]° offsets from the launcher’s moving trajectory separated by cardinal (blue) and oblique (red) launch direction. Each panel corresponds to a different binning of +/-[2.5, 5.0, 7.5]° to classify launches as cardinal and oblique in panels A, B and C, respectively. **D-F**. Causal impressions were on average higher in oblique launches than in cardinal launches, especially when +/- 5.0° and +/- 7.5° bins were applied for the cardinal / oblique categorisation. The bottom row shows the mean causal impression (dot) with the 95% confidence interval (bar) for cardinal (blue) and oblique (red) launches with individual participant data as grey lines in between. In most participants, there was a tendency to increase the causal impression for oblique launch directions.

When regarding the means with 95% confidence intervals in Figure 4 D-F it seems that oblique moving trajectories of the launcher resulted in stronger causal impressions (a stronger impression that the launcher’s and the target’s movement are connected) than cardinal moving trajectories of the launcher. The strength of that cardinal vs. oblique difference depended on the choice of binning of the launcher’s moving trajectory (two-sided Wilcoxon sign-rank test, *p1=0.18*, *p2=0.01*, *p3=0.04* for +/- 2.5°, 5.0°, 7.5°, respectively). The exact role of the dichotomised launcher’s movement direction is, however, unclear: while it indeed seems that oblique launcher moving trajectories on average yielded stronger causal impressions than cardinal launcher moving trajectories (Figure 4 D-F), the direction of the relationship is less clear for higher deviations (30° and 60°) from the launcher’s moving trajectory (x-axis in Figure 4 A-C). From this qualitative piece of evidence, it seems possible that the exact angle offset of the target’s relative trajectory could change the relationship between sensory uncertainty around the launcher’s trajectory and causal impressions.

The idea that the launcher movement and the target movement jointly influence causality ratings is aligned with a perspective that centres on the visual uncertainty around moving trajectories: in a full launching display, there is not a single movement trajectory but instead uncertainty around both the launcher trajectory (pre-collision) and the target trajectory (post-collision) to consider. Those two uncertainties are in our case naturally connected; for a cardinal launcher movement the target becomes more oblique, the stronger it disrupts the continuation of the launcher’s movement (higher angle offset). If the launcher ball instead was originally moving in an oblique direction, a higher angle offset in the target’s trajectory means that the target movement became more cardinal.

Given the inherent subjectivity of the causality judgment task, it is equally possible that strong disruptions of line continuation shift an observer’s criterion for attributing causality. Our current analysis does not allow us to discern between these two options (visual uncertainty vs. subjectivity), and they likely contribute together to a causal impression. To better understand how visual uncertainties for different movement directions may be linked to causal impressions, we further examined the effect of the launcher moving direction and the target moving direction. Since the evidence for cardinal / oblique contrasts depended on the choice of binning and we have no a priori reason to prefer one binning scheme over another, here we use the +/- 5° binning (producing the strongest cardinal / oblique contrast in the launcher moving angle) to also compute the mean cardinal / oblique contrast in the target’s moving trajectory in **Figure *5***. Note, that **Figure 5** C is equivalent to Figure 4 E and that Figure 5 A was created from the data underlying Figure 4 B.

**Figure 5.**
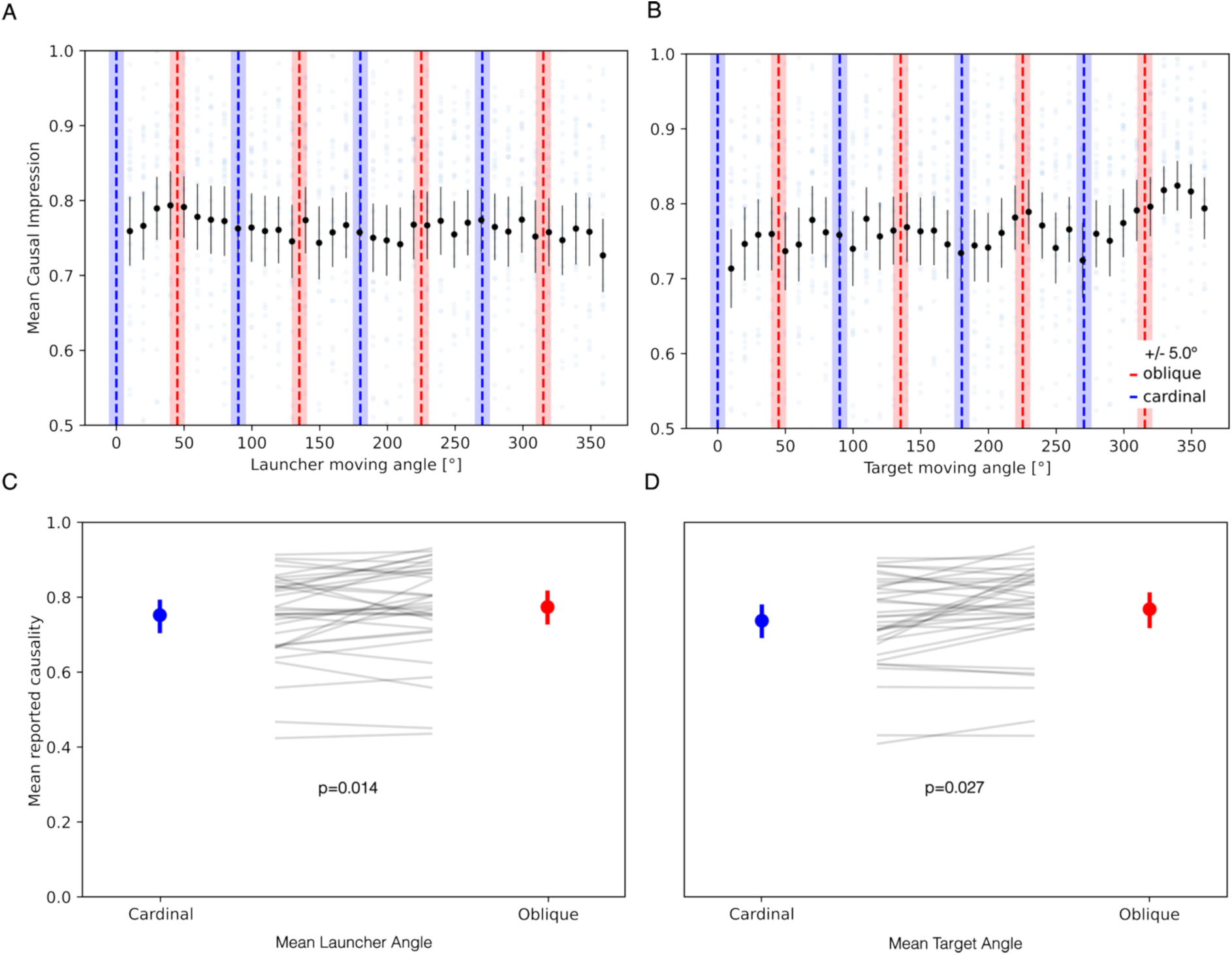
Mean causal impressions systematically depend on the launcher’s and target’s movement direction. **A.** Mean reported causality slightly fluctuates depending on the launcher’s moving angle. Faint grey dots show individual participant means and black dots show the mean of participant means per +/ -5° bins over different launcher movement directions. **B.** Same as A for the target movement directions. **C.** Causal impressions were overall increased in oblique (red) launcher movement angles compared to cardinal (blue) launcher movement angles. Means are depicted as dots with bars indicating 95% confidence intervals. Single-participant data is visualised as faint grey lines. Note, that this panel is equivalent to Figure 4E. **D.** Causal impressions were also increased in oblique (red) vs. cardinal (blue) target movement directions. Means are depicted as dots with bars indicating 95% confidence intervals. Single-participant data is visualised as faint grey lines.

Similar as shown above for the dichotomised launcher moving trajectory (Figure 4 D-F), launching events with an oblique target moving trajectory are on average judged as more causal than for a cardinal target moving trajectory (**Figure 5** D). For both oblique launcher movement and oblique target movement the causal impressions were on average increased (launcher oblique *M=0.77*, *SE=0.020* and target oblique *M=0.77*, *SE=0.020*) compared to cardinal movement (launcher cardinal *M=0.75*, *SE=0.020*, *p=0.014* and target cardinal *M=0.73*, *SE=0.020*, *p=0.027*, two-sided Wilcoxon sign-rank test at +/- 5.0° bins). Interestingly, the absolute target trajectory qualitatively leads to a consistent periodicity in the mean causal impressions (**Figure 5** B); that periodicity induced by the movement direction is less obvious in mean causal impressions plotted against the launcher’s moving angle (**Figure *5***A). Note, that faint grey dots in the background show individual participant means per bin and that the cropping of the y-axis to the range between 0.5 and 1.0 resulted in the exclusion of 82 (out of 1200) individual dots from the plot in Figure 5 A and B.

Overall, it is possible that the observed cardinal / oblique crossover at a relative target angle manipulation of 30° (see Figure 4) is indeed a consequence of the sensory uncertainty induced by the target moving angle and the data illustrated in Figure 5 could be grounds to explore that assumption further in future work. Note, that like for the launcher’s trajectory, the choice of binning of the target’s moving trajectory can change the strength of the cardinal / oblique contrast in causality ratings (two-sided Wilcoxon sign-rank test, *p1=0.027*, *p2=0.027*, *p3=0.0001* for +/- 2.5°, 5.0°, 7.5°, respectively).

To explore whether visual uncertainties interact with causal impressions under stronger and weaker violations of contiguity, we pooled trials by dichotomised launcher moving angle (cardinal / oblique) and by qualitative target movement continuation given the launcher’s trajectory (causal / not causal). Whether a collision was perceived as a high vs. low violation of continuity was determined post-hoc from the participants’ reported mean causal impressions applying a threshold of 0.5. Mean causal impressions dropped below that threshold at a relative target trajectory of 30° (Causal impression was *M=0.44*, *SD=0.25).* Using that cutoff criterion, launches with a target angle manipulation < 30° deviation from the launcher’s trajectory were labelled as high continuity trials; the rest of the trials (target angle manipulations of 30° and 60°) were considered to have low continuity (strong disruption of launcher moving direction).

In high movement continuity trials, we again find a systematic increase for causal impressions in oblique launcher moving directions compared to cardinal launcher moving directions (Figure 6C-D, cardinal *M=0.86*, *SE=0.021* (blue), oblique *M=0.87*, *SE=0.022* (red), two-sided Wilcoxon sign-rank test, *p=0.009* for +/- 5.0° binning). There seems to be no such tendency for a cardinal / oblique difference in low movement continuity trials (cardinal *M=0.30*, *SE=0.029* (blue), oblique *M=0.30*, *SE=0.029* (red), two-sided Wilcoxon sign-rank test, *p=0.66* for +/- 5.0° binning), implying that the relationship between sensory uncertainty and causality can change depending on the strength of violation of contiguity.

**Figure 6.**
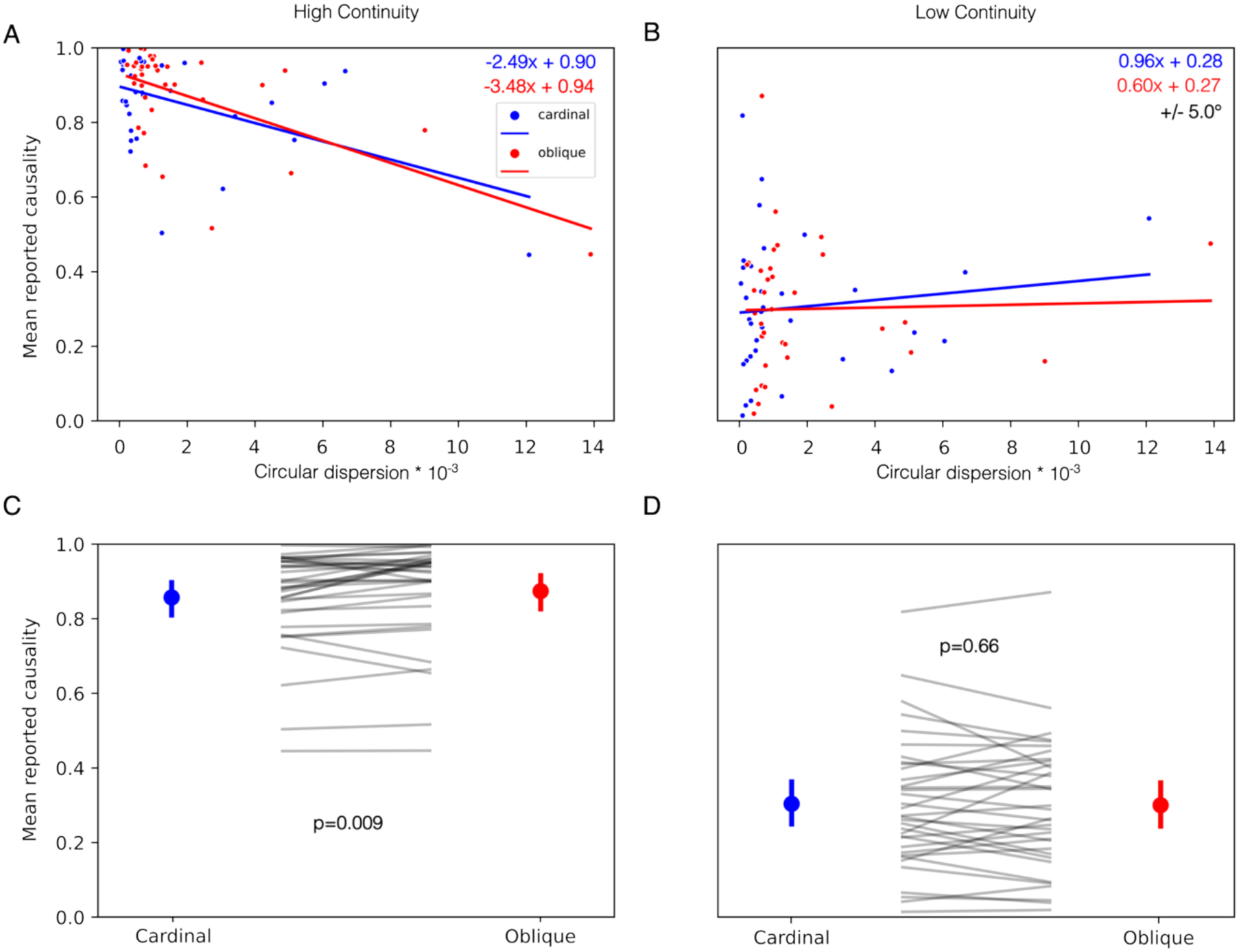
The relationship between sensory uncertainty and magnitude of the causal impression depends on the contiguity of the launcher’s and the target’s moving directions. **A.** The magnitude of individuals’ mean causal impressions correlates negatively with their respective mean uncertainty in cardinal (blue) and oblique launches (red) in high contiguity trials (< 30° offset between the target’s and the launcher’s trajectory). Oblique launches induced stronger causal impressions in high contiguity scenarios than cardinal launches (compare panel C). **B.** Same as A but the correlation is absent for low contiguity trials (30° and 60° offset from the launcher’s moving trajectory) in cardinal (blue) and oblique launches (red). Given the low values for the spearman correlation it is difficult to make a statement about the relationship between sensory uncertainty and cardinal / oblique differences in low contiguity trials (compare panel D). **C.** Oblique launches increase the magnitude of the causal impression in high contiguity trials. Dots depict the mean causal impression of the population and error bars depict the 95% confidence interval. **D.** The mean causal impression in cardinal vs. oblique trials with low movement continuity (blue and red dots with 95% confidence intervals, respectively) are not significantly different (but note that there were less underlying trials where the movement direction was disrupted than the number of trials where the movement was continued in panels A, C).

We furthermore observed that participants who were more uncertain in the trajectory prediction task tended to report weaker causal impressions in high continuity trials. Specifically, we observed a weak negative relationship between the circular dispersion of the predictions and an individual’s mean causal impression here (Figure 6A, cardinal spearman correlation coefficient of -0.29 with *p=0.090*, oblique spearman correlation coefficient of -0.43 with *p=0.010*). This result seems contradictory with our earlier findings: Figure 3D shows that on average, uncertainty around predictions was highest in oblique launches and Figure 4 implies that causal impressions are on average stronger for oblique launches. If sensory uncertainty and causality judgments share a straight-forward relationship we might generally expect stronger causal impressions given larger uncertainty (positive slopes) in Figure 6A-B (the opposite of the regression line for high continuity trials).

In low continuity trials, the correlation between sensory uncertainty and causal impressions seems to be non-existent or reversed compared to high continuity trials (**Figure 6** B, cardinal spearman correlation coefficient of 0.074 with *p=0.674*, oblique spearman correlation coefficient of 0.079 with *p=0.651*). Increased uncertainty in low continuity trials seems to increase the magnitude of the causal impression most prominently in cardinal launches, expressed by a steeper slope than in oblique launches. Note however, that our analysis does not show that cardinal vs. oblique launches change the magnitude of the causal impression in low continuity trials with disrupted moving directions on average (see Figure 6 D).

To understand whether differences between cardinal and oblique slopes in Figure 6 A and B were quantitatively supported, we performed paired permutation tests on the slopes of the linear fit for high continuity and low continuity trials, respectively. Specifically, we randomly shuffled the launch direction label on the data points in Figure 6 A, fit a linear regression to each nominal cardinal and oblique dataset, took the difference between the regression slopes, and then repeated this procedure 9,999 times to create a sampling distribution for the regression slope differences under a null hypothesis. The same procedure was repeated for Figure 6 B. The reported p-values are the two-tailed percentiles for the empirical slope difference.

The test confirmed that the observed cardinal and oblique differences were meaningful in high continuity trials (*p=0.0002* for +/- 5.0° bins) but were plausible under the null for low continuity trials (*p=0.52* for +/- 5.0° bins). The differences between high continuity and low continuity trials here could mean that causality judgments were more driven by sensory uncertainty around the predictions if the launcher’s and target’s moving direction were aligned. The more contradictory the target moving angle and the launcher moving angle, the less important is the visual uncertainty around the prediction. However, due to the imbalance of employed offsets, the underlying number of low continuity trials per participant was relatively small (15,660 vs. 3,500) leading to a noisy fit that has to be interpreted with caution; therefore, we cannot make a definite statement about whether continuity of launch trajectories would consistently change the slope of the regression line in a more representative dataset.

The correlation between sensory uncertainty and causal impressions (Figure 6A) could also be driven by outliers. Excluding two participants with very low causal impressions (< 0.6) in high continuity trials yielded different slope values of *M=-0.88* (cardinal) and *M=-1.79* (oblique) with lower spearman correlation coefficients (of -0.19 and -0.33 with *p=0.30* and *p=0.06* for cardinal and oblique, respectively). After the removal of these participants, the differences in corrected slopes of the correlation were no longer meaningful (pairwise permutation test, *p=0.52*). In that context, we report that there is a general inflation of very high mean causal impressions in our dataset, meaning values are generally skewed towards 1.0 (also visible in Figure 4 A-C). It is possible that correlations between mean uncertainties and causal impressions are masked by that inflation of responses because higher mean values of causality ratings are generally closely spaced together across a small range of the causality rating scale.

## Discussion

Our research question centred on how predictive uncertainty around moving trajectories influences causality judgments and consequently whether causal impressions arise from a comparison between predicted and observed post-collision motion. To empirically capture this predictive component, we measured both the directional bias, and the dispersion of participants’ trajectory estimates and treated circular dispersion as an index of sensory uncertainty. This allowed us to test, within the same observers, how directional anisotropies in prediction precision relate to the strength of perceived causality.

Across tasks, prediction uncertainty varied systematically with launch direction, showing the expected overall increase for oblique compared to cardinal trajectories (Duan et al., 2025), despite notable directional idiosyncrasies (Figure 3). Causal impressions likewise depended on both launch direction (Figure 4 and Figure 5) and the plausibility (high/low continuity) of the post-collision trajectory: oblique launches were associated with slightly higher causal impressions in plausible displays with high continuity, whereas this relation diminished or reversed for strongly implausible trajectories with low continuity (Figure 6). Together, these findings indicate that the mapping of physical plausibility onto a causal impression is not straight-forward: observers could base their individual weightings of predicted vs. observed motion on the general plausibility of the event in terms of moving trajectory.

In the prediction data, our results show an anisotropic bias in the predictions of moving trajectory (Figure 3B-D), where accuracy in the predictions systematically depends on the direction of the launch. Complementing our earlier report (Duan et al., 2025), we here provide an empirical measurement of uncertainty around sensory predictions. Directional dependencies of visual precision are consistent with the broader literature on anisotropies in spatial vision and motion processing, where cardinal orientations typically support higher perceptual discriminability than oblique ones (e. g. Gros et al., 1998). However, the changes in uncertainty due to our manipulation of launch direction (cardinal/oblique) are not as straight-forward as initially anticipated. For instance, Figure 3C clearly shows higher uncertainty for 270° (downwards movements) compared to the oblique orientations of 135° and 315°, although we would expect higher uncertainties at those oblique orientations. Nevertheless, we find that on average, participants had larger uncertainty in oblique launches given the difference in means of the circular dispersion for cardinal vs. oblique launch orientations (Figure 3D).

One would expect discriminability to be best at the horizontal moving directions and to be better at horizontal and vertical (cardinal) lines than in oblique lines (A-Izzeddin et al., 2024; Girshick et al., 2011; Gros et al., 1998). Except for 270° that seems to be true for our data (Figure 3). Interestingly, 270° describes a downwards launcher moving direction. Given that participants did not receive information on the context, their beliefs about the circumstances of the collision remain unclear: participants possibly combine their visual uncertainty around the downward trajectory with their individual belief of a gravitational bias (Hubbard, 2020). Due to the high level of visual abstraction in our collision (2D discs without surface properties, no friction, no velocity change), it is unknown whether observers had such biases and how that affected their prediction accuracy. In addition, observers’ sensory uncertainty around orientation can be different depending on the individual and context of a stimulus (A-Izzeddin et al., 2024; Van Geert et al., 2022).

We confirm that subjective causal impressions were qualitatively aligned with real-life implications of Newtonian mechanics (Figure 4), consistent with earlier demonstrations that observers judge collisions as more causal when post-collision motion adheres to physical contingencies (e.g., Michotte, 1963; White, 2012; Wende et al., 2013). This pattern aligns with accounts proposing that observers rely on a noisy mental simulation to infer causality from object interactions (Gerstenberg et al., 2012, 2021; Sanborn et al., 2013). At the same time, our results are compatible with evidence that causal impressions rely on early perceptual mechanisms tuned to motion direction and stimulus configuration (Kominsky & Scholl, 2020; Rolfs et al., 2013; Ohl & Rolfs, 2023). Our empirical results here therefore replicate key features of our earlier work (Duan et al., 2025) using launch directions drawn from the full direction circle on each trial, making it unlikely that these results were driven by participants using separate report strategies for horizontal and oblique launch categories.

It is essential to note that the target moving angle seemed to influence the mean causal impression (Figure 5). This observation fits with prior reports that judgments of causality can be modulated by stimulus configuration, contextual cues, the spatial trajectory of the moving objects and the subjective impression of a launch (Meding et al., 2020; Vicovaro, 2018; Vicovaro et al., 2020), beyond the relative offset alone (White, 2012). While we cannot grasp the comprehensive implications of the target’s moving angle in this work, investigating how uncertainties around the launcher’s and the target’s moving trajectory interact provides an interesting avenue for future research.

Our data are also consistent with the possibility that the offset from a plausible trajectory does not simply monotonically influence causal impressions but alters the relationship between causal impressions and sensory uncertainty (Figure 6); a similar perspective has already been proposed for objects with specific vs. objects with abstract material properties (Vicovaro, 2018). In our account, plausibility (here manipulated using implausible target moving angles) could change how sensory evidence is weighted by the observer: in low continuity displays participants may judge causality more subjectively and dismiss the physics. These complexities resonate with previous work showing that causal impressions are shaped not only by spatial–temporal contingencies but also by contextual interpretation (Sommer et al., 2025) and sensory realism (Meding et al., 2020). For instance, it has been demonstrated that the same manipulations of movement trajectory and temporal delay can have opposite effects on causal judgments depending on whether observers interpret an event as physical or social, and that no unitary causal-processing network is engaged across contexts (Blos et al., 2012). This may be seen as evidence that observers flexibly draw on different evaluative strategies, consistent with our interpretation that causal ratings in our task could reflect a mixture of predictive and postdictive mechanisms.

Overall, we cannot fully dismiss the idea that the degree of launcher movement continuation or disruption by the target movement induces the cardinal / oblique differences influencing causality judgments. The fact that the relationship between sensory uncertainty and causality judgments is influenced by outliers (Figure 6) may be seen as confirmation that the relationship between perceptual uncertainty and subjective causality ratings is difficult to capture. On a similar note, we qualitatively observed that similar uncertainty scores in Figure 6 A-B, especially around 0-2 * 10^-3^ circular dispersion units, can lead to very different causal impressions (indicated by the spread of data points in y-direction). That finding further supports the idea that sensory uncertainty and causality judgments share a complex relationship that might not be easy to quantify.

Another possibility for the observed variability in our data is that the observer has no direct access to the predicted post-collision moving trajectory because the pre- and post-collision evidence become perceptually fused into one. That would mean that the observer performs a Gestalt-like integration of pre- and post-collision trajectories into a single angle, without a proper retention of the pre-collision trajectory and the respective prediction of motion. Such a mechanism has already been proposed for launcher and target speed perception in launching displays (Kerzel et al., 2000; Vicovaro et al., 2020) and is consistent with the idea of discrete perception (i. e. Herzog et al., 2020). Future research and different studies focusing on empirical assessments of visual uncertainty are necessary to examine the role of predictive and postdictive mechanisms in attributing causality.

## Conclusion

In this work, we provide an empirical assessment of sensory predictions in launching displays by combining a trajectory-prediction task with causality ratings from the same observers. This approach allowed us to quantify direction-dependent uncertainty in the sensory predictions that precede a causal impression and to examine whether this anisotropic uncertainty relates to mean reported causality. While causality ratings were broadly aligned with the physical plausibility of the post-collision trajectory, the relationship between prediction uncertainty, launch direction, and perceived causality was more complex than expected. Instead of relying on a single predictive strategy, observers could be flexibly weighting internal predictions, perceptual cues, and prior expectations about temporal contiguity depending on the plausibility of the collision. This flexibility furthermore implies that causal attributions cannot be reflected in a feed-forward fashion but can be shaped retrospectively by perceptual organization and the subjective impression of a launch. Together, our findings highlight the importance of characterising how predictive and postdictive mechanisms jointly contribute to causal perception and underscore the need for models that explicitly incorporate direction-dependent uncertainty in both pre- and post-collision motion processing.

## Acknowledgments

We thank Lukas Maninger for useful comments on the manuscript and code review. We thank Dr. Yunyan Duan for early feedback on the concept and for the code implementation of the pooltool physics engine.

## Contributions

The experiments presented in this work were conceptualised by LEK and TW in collaboration with FT, CS and BS and feedback from Yunyan Duan (YD). Software to conduct the experiment was implemented by LEK and YD and expanded by Lukas Rosenberger (LR). LR recruited participants and performed the data acquisition. Resources to conduct the experiments were provided by TW. Software and methodology for analysis was developed and implemented by LEK, LR and TW with feedback from FT and code review from Lukas Maninger (LM). Data was curated by LEK and LR. LEK, LR, TW and FT were involved in developing the visualisation of the data. LEK had the administrative oversight of the project under the supervision of TW and in collaboration with CS and BS. LEK took the lead in writing the manuscript with input from TW and feedback on the discussion from LM. LEK, FT, CS, BS and TW reviewed the manuscript critically and were involved in editing it.

## Data and code availability

All anonymised data and code used in the analysis along with the experimental scripts and demo videos are available at: https://doi.org/10.5281/zenodo.18511298.

